# Cryogenic light microscopy with Ångstrom precision deciphers structural conformations of PIEZO1

**DOI:** 10.1101/2024.12.22.629944

**Authors:** Hisham Mazal, Alexandra Schambony, Vahid Sandoghdar

**Author notes:** Correspondence to: Vahid Sandoghdar.

## Abstract

Despite the impressive progress in molecular biochemistry and biophysics, many questions regarding the conformational states of large (transmembrane) protein complexes persist. In the case of the PIEZO protein, investigations by cryogenic electron microscopy (Cryo-EM) and atomic force microscopy (AFM) have established a symmetric trimer structure with three long-bladed domains in a propeller-like configuration. A transition of PIEZO protein from curved to flat conformation is hypothesized to actuate closed and open channels for the flow of ions. However, conclusive high-resolution data on the molecular organization of PIEZO in its native form are lacking. To address this shortcoming, we exploit single-particle cryogenic light microscopy (spCryo-LM) to decipher the conformational states of the mouse PIEZO1 protein (mPIEZO1) in the cell membrane. Here, we implement a high-vacuum cryogenic shuttle to transfer shock-frozen unroofed cell membranes in and out of a cryostat for super-resolution microscopy at liquid helium temperature. By localizing fluorescent labels placed at the extremities of the three blades with Ångstrom precision, we ascertain three configurations of the protein with radii of 6, 12, and 20 nm as projected onto the membrane plane. Our data suggest that in the smallest configuration, the blades form a nano-dome structure that is more strongly curved than previously observed and predicted by AlphaFold-3. In the largest conformation, we believe the structure must fully unbend in an anticlockwise manner to form a flat extended state. We attribute the 12 nm conformation, the most frequently occupied state, to an intermediate state and discuss our results in the context of the findings from other groups. Combination of spCryo-LM and Cryo-EM measurements together with *in situ* photothermal stimulation promises to provide quantitative insight into the interplay between structure and function of PIEZO and other biomolecular complexes in their native environments.

---

PIEZO proteins are large transmembrane mechanosensitive ion channels that are involved in the translation of mechanical forces to biological signals and steer the regulation of various physiological processes^1^. The PIEZO1 protein is composed of 2547 amino acids (**Fig. 1a-b**) and is predicted to form 38 transmembrane alpha helices (TM)^2,3^. The structure was investigated by Cryo-EM^3-6^ and AFM^7^, and it was found to consist of a symmetric trimer (**Fig. 1b**), where the C-terminal domain forms the core of the channel (**Fig. 1a-b**). The N-terminal domain in each protomer is composed of 36 transmembrane alpha helices, forming a long blade-like structure whose end points lie on a circle of radius *r* from the core (**Fig. 1a-d**). Previous studies have revealed that the blade domains curve out the membrane plane into a nano-dome structure^3,8,9^ (**see Fig. 1d**), inspiring the hypothesis that the channel is closed in the curved state, and it opens for ion flow when the structure flattens in response to an induced membrane tension^3^. In other words, curvature plays a central role in mechanosensing^9-13^.

**Fig. 1.**
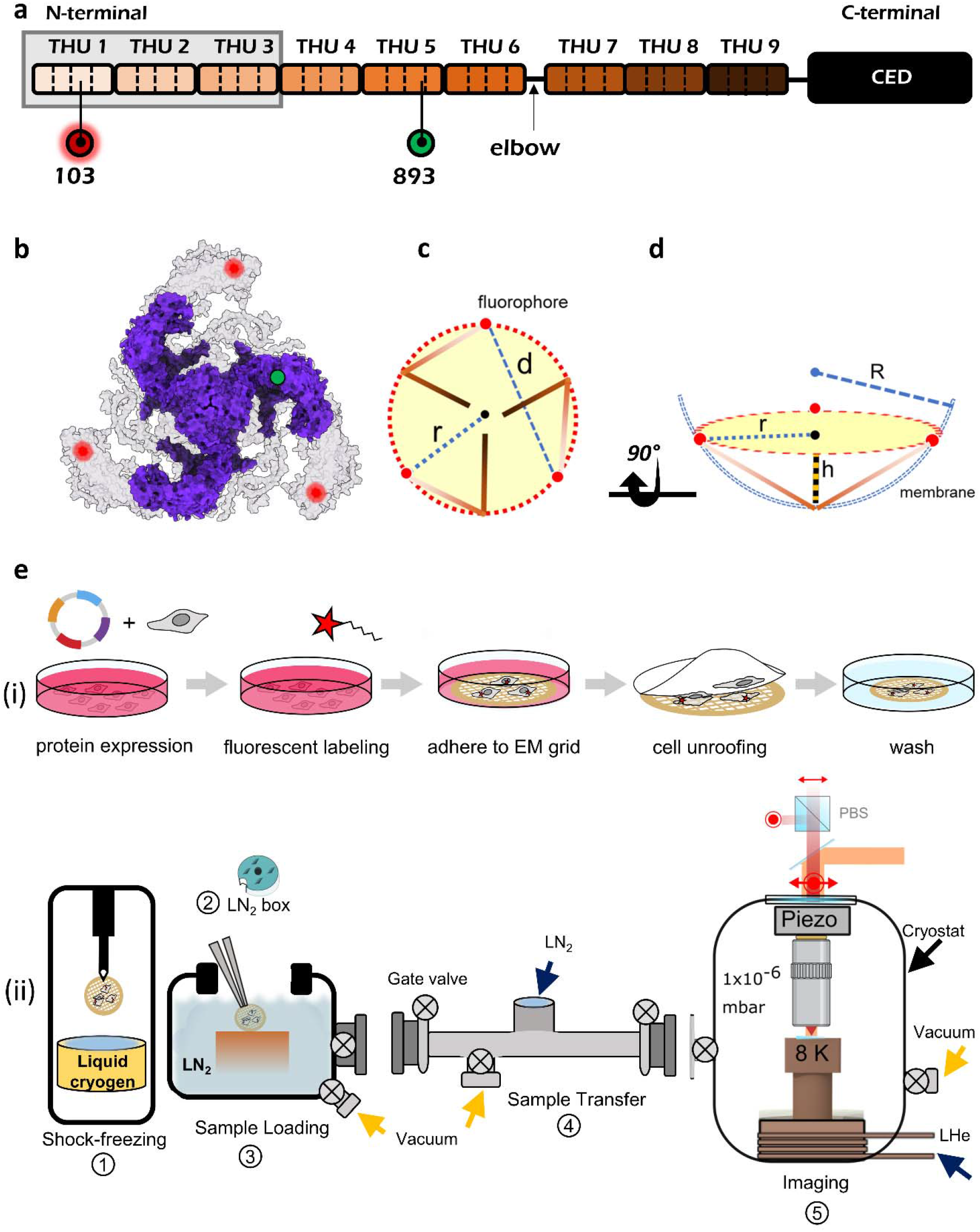
Imaging mPIEZO1 protein in near-native cell membrane. **a**, Domain organization of monomer mPIEZO1 protein composed of 9 transmembrane alpha-helical units (THU), colored in a gradient of brown. Each unit is a bundle of 4 alpha helices, separated by black dashed lines. The C-terminal extracellular domain (CED) which forms the core of the channel is shown in black. Positions 103 and 893 mark the labelling positions mentioned in the text. The arrow indicates the elbow region at which the blade structure curves. **b**, Cryo-EM structure of mPIEZO1 (PDB: 63BR) in purple docked into the AlphaFold-2 predicted structure in gray (E2JF22). Red and green spots show the exact location of labelling position amino acids 103 and 893. **c**, Schematic top view of the nano-dome structure. Brown lines are the blade domains. Dashed and dotted blue lines show the inter-blade distance (*d*) and in-plane radius (*r*). d, Side view of (c), illustrating the dome model of PIEZO protein as explained in Ref.^12^. The dashed blue lines depict the curved membrane with radius *R*. **e**, (i) Pipeline of sample preparation. (ii) Schematic view of the five major steps in the workflow. (1) Sample is plunge frozen in cryogenic liquid in a dedicated commercial instrument. (2) Sample is placed and stored in a liquid nitrogen (LN_2_) TEM grid box. (3) The grid box is transferred to a chamber, where it is loaded onto a special sample cartridge inside a small LN_2_ dewar. Then the chamber is put under vacuum. (4) A transfer chamber under vacuum is used to move the sample to the measurement chamber. (5) Cryogenic optical microscope operating at liquid helium temperature.

Despite these advances, our understanding of the structure and function of PIEZO proteins remains incomplete owing to the complexity of this protein and various experimental challenges. In particular, a significant part of the blade domain remains unresolved in Cryo-EM measurements (**Fig. 1a-b**). Importantly, much of the existing data have been obtained from isolated proteins reconstituted in detergent or synthetic membranes^3,4,6-8^. Considering that the details of lipid composition are believed to regulate the PIEZO activity in cells^1,14^, high-resolution measurements on native cell membranes become indispensable. However, previous efforts toward deciphering the blade structure were realized in chemically fixated plasma membranes of cells and lacked sufficient spatial resolution and statistics^15,16^. We now exploit a new imaging strategy in which we unroof cells, plunge freeze them, and then transfer them with a vacuum tight cryogenic shuttle to a cryogenic super-resolution fluorescence microscope, where single PIEZO proteins are imaged with Ångstrom precision. In the discussion section of this article, we put our findings in the context of the previous state-of-the-art investigations^3-9,15-17^.

## Optical imaging at Ångstrom precision of mouse PIEZO1 protein in vitrified cell membrane

Ångstrom resolution in structural studies requires the elimination of all thermal and functional motion that would smear spatial information. Room-temperature microscopy achieves this via chemical fixation^18,19^, which generally includes harmful substances to the cells and might result in artifacts and distortions of the biomolecular structure^20-22^. To circumvent this complication, electron microscopists have developed vitrification of biological samples via shock-freezing^23^, where water molecules are nearly instantaneously frozen into an amorphous ice state (vitreous ice), thus preserving the hydrated environment^22,24,25^. This method has recently been also employed to correlative Cryo-EM and Cryo-LM studies, where the latter served as annotation while the former delivered high-resolution images^26-30^. In this work, we use shock-frozen samples to directly obtain cryogenic fluorescence images of individual biomolecules with Ångstrom precision.

A few years ago, we showed that fluorescence microscopy at liquid helium temperatures brings about the decisive advantage that photochemistry and photobleaching are reduced, leading to orders of magnitude more collected photons per fluorophore^30^. Here, we exploited the stochastic photoblinking of individual fluorophores and resolved several fluorescent dye labels on a single protein by sequential localization of each individual dye with few Ångstrom precision^31,32^. Moreover, we could decipher the 3D orientation of each structure in an analogous fashion to single-particle cryogenic electron microscopy (spCryo-EM), but the demonstrations used protein structures embedded in a polymer^31,33^. To adapt our approach to near-native imaging of transmembrane proteins, we constructed a shuttle system that allows us to transfer vitrified samples in and out of the cryogenic optical microscope with negligible condensation and devitrification^34^. **Figure 1e** sketches the principal workflow^34^.

We image three fluorescent dyes placed on mPIEZO1 proteins. In order to probe the blade conformational state, we used a previously generated DNA construct^15^ that incorporates an unnatural amino acid trans-Cyclooct-2-en–L–Lysine (TCO*K) at position 103, close to edge of the unresolved part of the blade (**Fig. 1a,b**). The protein construct was expressed in COS7 cells since these cells adhere strongly and spread on the substrate. Following protein expression, the cells were labelled specifically at position 103 using the small organic fluorophore Pyrimidyl-Tetrazine-AF647. As plunge freezing is most effective for sample thickness less than 1 µm^23,25^, we followed a protocol^35,36^ to unroof the cells, leaving only their basal membranes intact on the TEM grid (see **Fig. 1e(i)**). This technique generally keeps the cell cytoskeleton connected to the cell membrane^16,36-38^ and has been shown to maintain Ca^+2^-dependent exocytosis activity^36^. Indeed, by labeling the actin filaments of the unroofed cell, we confirmed their presence across the cell membrane. We, therefore, assume that the lateral membrane tension is preserved upon unroofing.

**Figure 2a** shows exemplary bright-field microscope images of several regions of a TEM grid carrying cells, and **Fig. 2b** displays a fluorescence image from part of a grid field. **In Fig. 2c**, we present a close-up image of a 15×15 μm^2^ region from (b), revealing a sparse distribution of fluorescent spots. The low density of PIEZO proteins in the membrane facilitates their distinction from each other within the diffraction limit. It, thus, follows that each spot, corresponding to a point-spread function (PSF) of the optical system represents up to three fluorophores. Photophysics characterization of PIEZO proteins labeled with Pyrimidyl-Tetrazine-AF647 showed fast blinking events, with a blinking off-on ratio of ∼5, and an average photon count rate of ∼5000/s, suitable for high-resolution imaging. As a validation check, we also imaged labelled non-transfected cells to assess the autofluorescence signal and unspecific binding of fluorescent molecules.

**Fig. 2.**
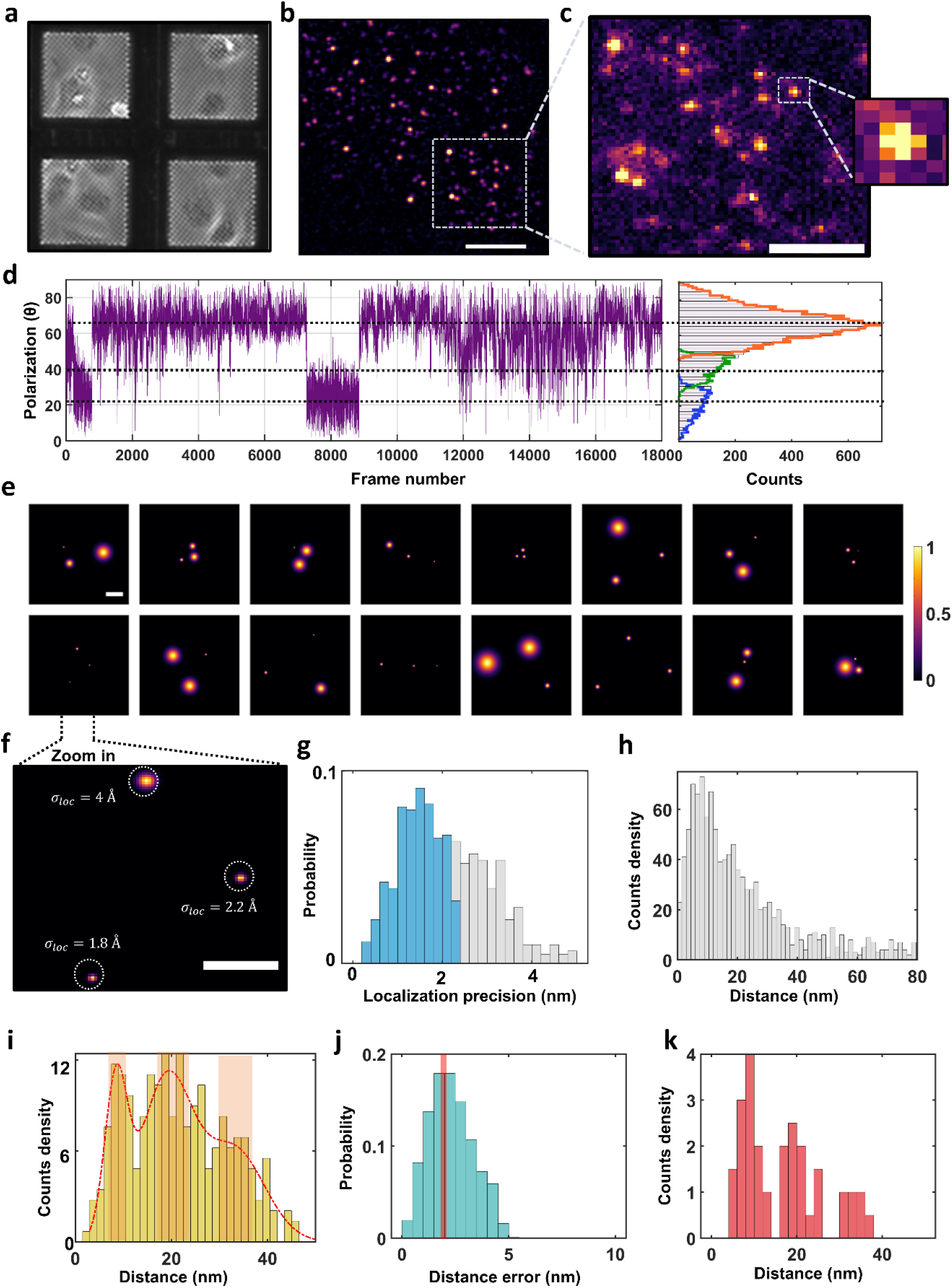
spCryo-LM imaging and analysis of mPIEZO1 protein. **a**, Bright-field image of COS7 cells adhered to UltraAUFoil-2C TEM grids. Each mesh square is 100×100 μm^2^. **b**, A fluorescence image taken by our cryogenic optical microscope from one of the mesh squares as demonstrated in (a). The image is median filtered with a kernel size of 5×5 pixels. Scale bar is 10 μm. **c**, Distribution of fluorescence spots across a grid region of 15×15 μm^2^ from (b), without any processing (raw image). Inset shows a close-up of one PSF with pixel size 227 nm. Scale bar is 5 μm. **d**, Exemplary polarization time trace extracted from one of the detected PSFs. We note that this trace is not a continuous time trace, but it rather represents the on-states of the fluorophores during the recording, i.e., off-times were omitted. The histogram in the right panel shows the polarization time trace decomposed to three states obtained from an unsupervised signal fitting approach^49^. **e**, 2D resolved images generated from particles with three polarization states reveal projections of different individual proteins in the sample plane. The spots represent the location of the fluorophores, and the width represents the uncertainty of their location. Color bar indicates the normalized probability of fluorophore position. **f**, Close-up of a particle indicated in (e) demonstrates Ångstrom localization precision. Scale bar in panels (e-f) is 5 nm. **g**, Histogram of localization precisions from ∼ 615 particles. The blue area indicates particles with localization precision better than ∼2 nm. **h**, Pairwise distance histogram. **i**, Histogram of maximum side lengths from each particle. Red curve shows a fit based on a Gaussian mixture model, yielding three components based on Akaike information criterion. The highlighted regions (light red) indicate the selected data for visualizing the aligned 2D images. **j**, Estimated error of the measured distance, using error propagation. The median value indicates an error of 2.3 nm. The red line marks the criterion to filter the maximum distance histogram in (i). **k**, Maximum distance histogram in (i) after filtering to include only distances with an error below ∼ 2.3 nm.

Given that in the frozen state the emission dipole moment of each dye molecule is randomly fixed in space, the three fluorophores on a protein can also be distinguished through polarization-selective detection^32,33^. In the current study, we chose PSFs that displayed three polarization states (see **Fig. 2d**), amounting to a total of N = 615 analyzed particles. **Figure 2e** shows a mosaic of 2D projections from 16 proteins. The size of each spot in these images signifies the precision with which it was localized. **Figure 2f** presents a close-up of the fluorophore positions from one protein, where all three were localized with Ångstrom precision. In **Fig. 2g**, we plot the distribution of the localization precision for the total population, with a median at 1.3 nm and an average precision of 1.4 nm per protein. Selection of cases with average localization precision better than ∼2 nm for further analysis reduced the number of particles to 378. We note that the localization precision and the statistics could be further enhanced by extending the sample imaging time since photobleaching is nearly suppressed.

The overall distribution of the distances *d* between neighboring fluorophores projected onto the imaging plane gives rise to the grey histogram in **Fig. 2h**. The 3D-to-2D projection smears the major side lengths of a triangle toward shorter distances. To ascertain whether the observed broad distance distribution arises from diverse triangles with unequal side lengths or from several subpopulations of equilateral triangles with different side lengths, we aligned all the projections about their centers of mass and performed in-plane rotations. The outcome yielded a symmetric Y-shaped pattern, providing strong evidence for symmetric equilateral conformations. However, classification of different subpopulations within the broad histogram of **Fig. 2h** is not straightforward.

To address this difficulty, we exploited the fact that for randomly oriented equilateral triangles, one of the projected side lengths always remains very close to the triangle side length: in the case of no localization error, the projection falls within a small range bounded by *d* and 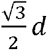^34^. **Figure 2i** displays the resulting histogram, revealing peaks at *d* ∼ 9, 19, and 34 nm, determined from a fit by a Gaussian mixture model. Thus, by taking the maximum side length from each measured protein, we minimize the smearing effect of 2D projections. We confirmed that the existence of three major peaks is not sensitive to binning parameters. Moreover, simulations indicate that the broadening of each subpopulation is, indeed, close to the localization precision limit. To estimate the error in the extracted values of *d*, we propagated the localization errors for each dye. As shown in **Fig. 2j**, the median of the resulting distribution corresponds to 2.3 nm. **Figure 2k** shows that if we filter the data to keep cases with distance errors < 2.3 nm and select polarization trajectories with signal-to-noise ratio larger than ∼3.5, the three peaks become resolved more clearly. **Figure 3a(i-iii)** visualizes the 2D projections corresponding to the three distance groups. The same results can also be obtained if we simply select the particles that belong to the central regions of each identified peak in **Fig. 2i** (marked by shaded bands).

**Fig. 3.**
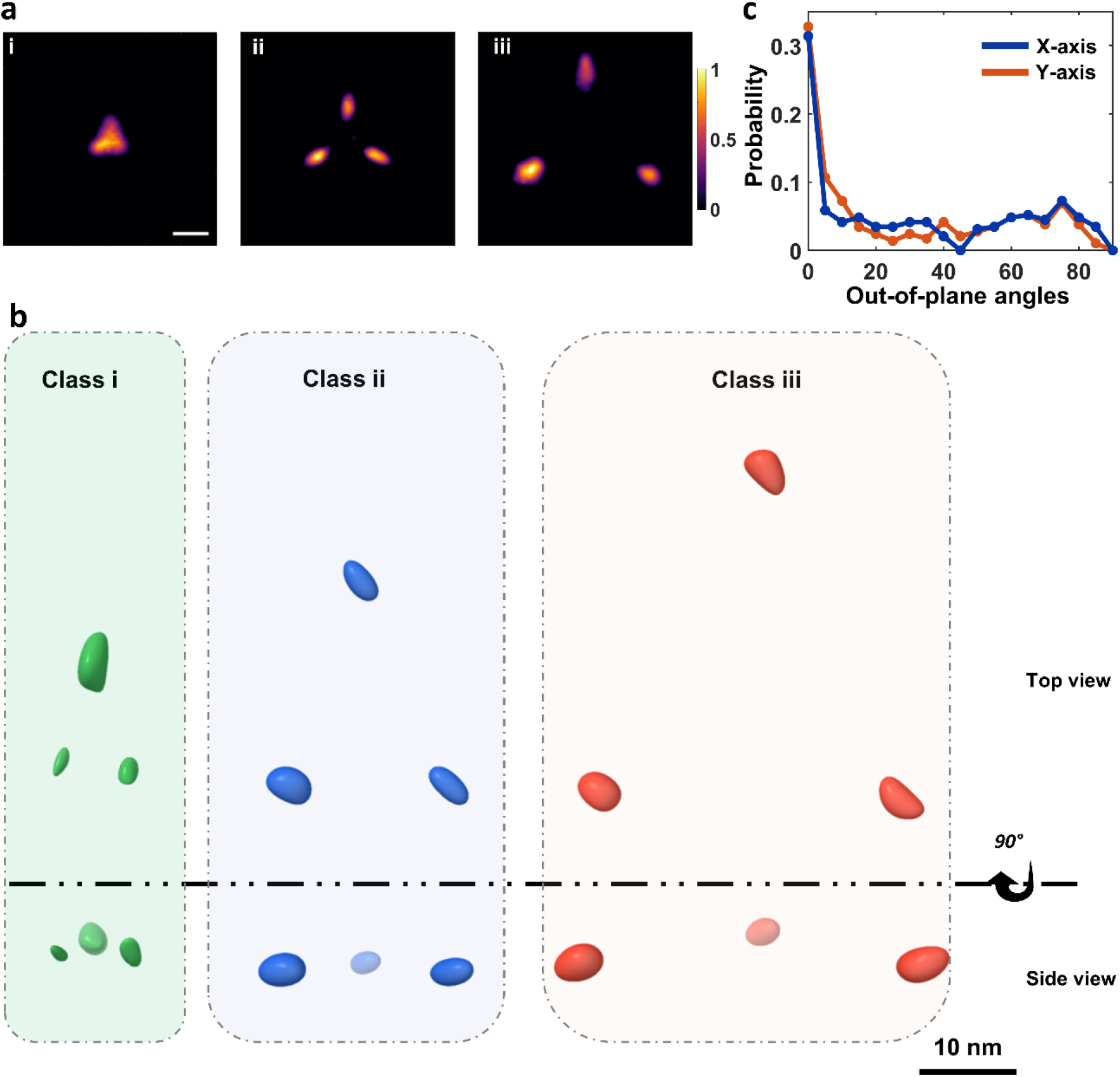
Dissecting the structural conformations of the mPIEZO1 protein. **a**, 2D projections after aligning all measured particles through rotations. Panels (i-iii) show the subpopulations for the three distances identified in Fig. 2: *d*= 9 (i), 19 (ii), 34 (iii) nm, establishing three classes of conformations. The final images were filtered against lower probability (below 0.5) of outlier projections. The smear of the localization spots is influenced by the localization precision and protein orientations (see panel c). Scale bar is 10 nm. **b**, Top-view and side-view projections of the 3D reconstructed volume of each class. Spheroids in green, blue and red indicate the localization uncertainties (see movies 4-6). **c**, Probability of out-of-plane orientation of the particles in the range of [0°-90°], as estimated based on simulated annealing algorithm. The blue and orange curve indicate the probability of the out-of-plane angles around the x-axis (pitch), and the y-axis (roll), respectively. As can be seen from the plot, the particles appear with the highest probability close to 0°, i.e., lying in the observation and membrane plane, while the probability of out-of-plane angles is lower.

Having established the three most probable equilateral triangle side lengths, next we employed a classification pipeline to include the data from all measured proteins in order to assess the fractional weight of each class. Here, we first generated 2D projection templates for equilateral triangles as model system. We classified each measured projection based on a template matching score between 0 and 1 to one of the side lengths of 9, 19 and 34 nm, yielding subpopulations of 21%, 51%, and 28%, respectively. Next, we used the classified projection to arrive at their 3D volume using a reconstruction algorithm^33,39^ with a precision of 4-8 Å as estimated from the Fourier shell correlation (FSC) analysis^40^. In **Fig. 3b**, we present the top (i) and side (ii) views of the projections from the resulting 3D reconstructions. **Figure 3c** shows that as expected for a transmembrane protein, proteins mostly lie in the membrane plane. Here, one nshould also bear in mind that a native plasma membrane is not fully flat but can contain regions of local curvature, where proteins are oriented out of the imaging plane.

## Discussion

A notable result of the first Cryo-EM studies of mPIEZO1 in detergent^3-6^ was that the blades project out of the plane. To investigate the mechanical interplay between the PIEZO protein and its surrounding membrane, several groups have reconstructed mPIEZO1 in synthetic liposomes^3,7-9^. These works have verified that the blade domain can curve the membrane into a dome-shaped structure when the protein is in its outside-in configuration (the C-terminal extracellular domain facing the inside of the liposome), acquiring an out-of-plane curvature *R* ∼ 10 nm (see **Fig. 1d**). In addition, measurements on the outside-out configuration could capture the flat conformation of the mPIEZO1 protein in synthetic liposomes with *r*=14 ^8^. As shown in **Fig. 1c,d**, top-view imaging measures distance *d* between certain blade locations and can yield *r*, whereas imaging membrane cross-sectional cuts can provide insight into the out-of-plane curvature of the protein and of the membrane. The quantities *r* and *R* can be connected via the Relation 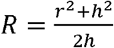, based on a spherical surface that reaches height *h*^12^.

AFM-based experiments could demonstrate the transition of mPIEZO1 from curved to flat state in real time when reconstituted in a supported lipid bilayer (SLB)^7^, although the direct link between the flat state and channel activity is still missing. While it is commonly assumed that the channel operates through a two-state mechanism (curved and flat), some studies have observed three states. For example, AFM-based experiments found two main populations with *r* ∼11 nm, 17 nm, and other minor population at *r* ∼22 nm^7^. Moreover, a very recent Cryo-EM study^17^ generated an activated state of mPIEZO1 via mutagenesis in a detergent environment and identified three dominant conformations with *r* ∼10 nm, 12 nm, and 12 nm and estimations of *R* ∼12 nm, 32 nm, and 110 nm, termed curved, intermediate, and fully flat, respectively.

Another study developed a model based on membrane elasticity theory to determine the shape of the mPEIZO1 dome in an asymptotically flat membrane and predicted *R* ∼42 ± 12 nm ^9,12^, which is significantly larger than the estimated value of *R* ∼ 10 nm in liposomes^7,8^. To examine the validity of the above-mentioned findings for proteins in native cell membrane, further questions have to be addressed: for example, are there other important contributions from cellular compartments such as the cell cytoskeleton, and how is the nano-dome structure affected by the complexity of the plasma membrane^11,13^. Several experimental efforts have, thus, investigated mPIEZO1 in natural cells., e.g., via STED (labeling position 893, see **Fig. 1a,b**) and MINFLUX (labeling position 103) microscopy techniques. However, these were done on chemically fixed samples with limited statistics and resolution. In both cases, only an ensemble average distance of *d* ∼ 25 nm was reported.

As the position of amino acid 103 on the blade is not resolved by Cryo-EM studies of mPIEZO1, we compare our data with the fully predicted structure model of mPIEZO1, generated by the artificial intelligence program AlphaFold-2 (AF2)^41^ and its most recent version AF3^42^. However, it should be kept in mind that as the trained data sets mainly stem from isolated protein models, exact structural prediction of protein structures in the native membrane environment, are not fully reliable. For example, AF2^41^ and AF3^42^ predicted different structural models of mPIEZO1.

To place our measured blade distances in the context of the previous reports^3,7,8,10,17^, we converted the measured blade distances into the in-plane radius *r* (**Fig. 1c**) following the relation 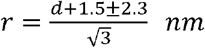, where *d* signifies the distance between the blades at position 103. The value 1.5 in the numerator is the approximated distance from position 103 to the edge of the blade estimated based on the AF2 model (E2JF22), and 2.3 nm signifies our experimental distance error (Fig. 2j). Following this procedure, we obtain *r* ∼ 6 (class i), 12 (class ii), 20 (class iii) ± 1.3 nm for the three observed configuration classes discussed above.

Our findings for the second class (*d*=19 ±2.3 nm, *r*=12 ±1.3 nm) are in excellent agreement with the predictions *d*=19.5 nm and *r* ∼12 nm of a structural model based on AF2 (E2JF22). For class i, however, our measured *d* and *r* values are considerably smaller than previous findings^3,7,8^. AF3 suggests that the blades are rotated further clockwise toward the center of the protein to yield *d* =13.5 nm and *r* ∼9 nm. Comparison of these data with our measurements *d* = 9 ± 2.3 nm, *r* = 6 ± 1.3 nm, suggests that in addition to the clockwise movement of the blades toward the center of the protein, they must also move out of the membrane plane. To examine this hypothesis, we considered the region between TM24 and TM25 (Fig.1a) as an “elbow” for the PIEZO1 blades^3,7^ and conducted rigid-body rotations of the blade arm out of the plane around the identified elbow region. This simple structural assumption can describe a small upward movement of the blades, ∼ 25°, yielding a highly curved state (see **Fig. 4a**).

**Fig. 4.**
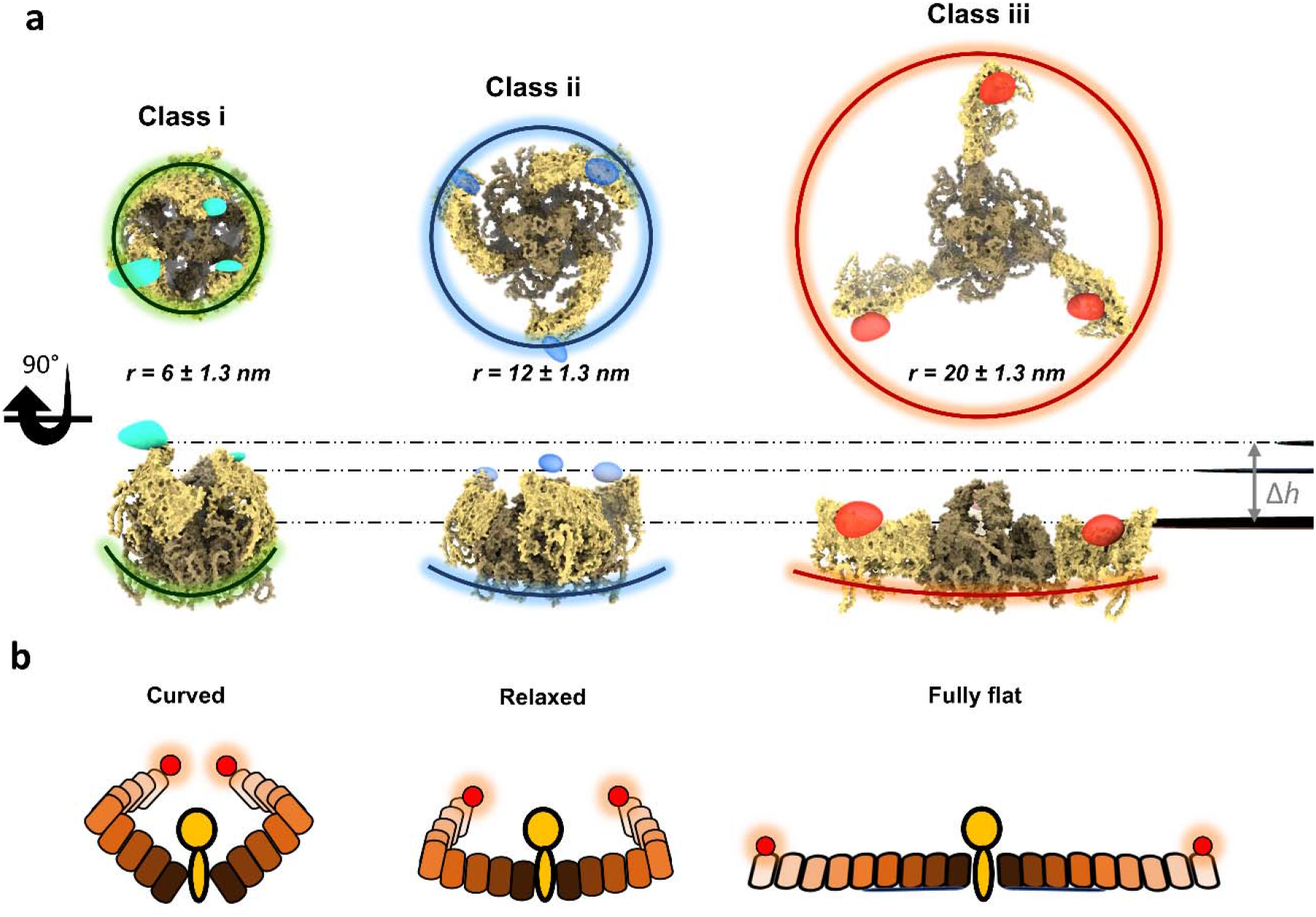
Structural modelling of the mPIEZO1 conformations. **a**, Structural models based on AF2 and AF3 obtained after rigid-body rotation (class i and class iii) and AF2 structure (class ii), which match our experimentally measured distances. Spheroids highlighted in green, blue and red indicate the 3D resolved volume using the classified 2D projections, for class i, ii and iii, respectively. Their position matches the structural model and the expected fluorophore position. Horizontal dashed lines indicate the planes of fluorophore locations. **b**, Schematics of the model deduced from our data, showing the curved, intermediate/relaxed and fully flat state of the blade domain. The difference in height of the ends of the blades between class iii and class i (see dashed lines in (a) amounts to ∼ 5.5 nm.

In the case of the largest distance between the blades in class iii, our data (*d* =34 ± 2.3 nm, *r* = 20±1.3 nm) indicate a larger structure than the flat conformation (PDB: 7WLU) of mPIEZO1 reconstituted in a lipid model system and resolved by Cryo-EM^8,17^, which yields *d* ∼ 27 nm and *r* ∼ 15 nm if we include the unresolved part of the blade . Here, we hypothesize that the blades move by an additional in-plane counterclockwise rotation around the previously mentioned elbow region, thus, generating a straighter shape and occupying a larger area than what has been reported previously. Indeed, by rotating the blade structure around the elbow, we found that an in-plane counterclockwise rotation of about ∼50⍰ very closely represents the measured distance of *d*∼ 34 nm, and *r* ∼ 20 nm, resulting in a fully flat and extended PIEZO state (see **Fig. 4a**).

We cannot directly assess the side-view radius of curvature *R* of the PIEZO-membrane system as discussed in previous works^9,17^. However, in **Fig. 4b** we examine the variations in height between the three scenarios based on the structural models that follows from our measurements (see also **Fig. 1d**). If we consider *h* = 0 for the (nearly fully) flat case of class iii, the height difference to the highly curved state of class i becomes Δ*h* ∼ 5.5 nm, in agreement with the range of *h* ∼ 0 - 5 nm proposed in the literature^7,17^. From our structural model and the maximum estimated height of class i, we then calculate *R* = 6 ± 2 nm, indicating mPIEZO1 in a highly curved membrane topography (**Fig. 4**). As there is some uncertainty regarding the h value of class ii, we calculated *R* over the estimated range of *h* = 0-5 nm, resulting in a median value of *R*=30 nm with 95% confidence interval of [26 – 35 nm] (**Fig. 4b**). Thus, we conclude that class ii conformation (51% of the population) must be a relaxed state with a more flat structure. Furthermore, we attribute class iii conformation to the nearly fully flat state of mPIEZO1 with completely unbent blade domain. We note that while *R* = ∞ for h=0, a slight deviation to h=1 nm would imply *R* ∼ 200 nm.

## Conclusions

Ångstrom precision in single-particle cryogenic light microscopy (spCryo-LM) allowed us to reveal three major conformational states in mPIEZO1 protein in the membranes of unroofed cells. To achieve these results, we introduced an experimental approach to circumvent the use of *in vitro* purification methods and chemical fixation, which have commonly been employed in high-resolution studies^18,19^. Our findings are in general agreement with previous results from Cryo-EM^3,8,17^, AFM^7^, and the structural prediction based on membrane elasticity theory^9,12^, but we provide direct and quantitative evidence for two extreme scenarios of the blade expansion and curvature, which were not clearly resolved by other methods. We believe a key advantage in our approach is the high signal-to-noise ratio of the localization procedure from each individual measured protein, thus, reducing the need for averaging processes that are currently necessary in Cryo-EM analyses. This allows us to sort and classify particles more accurately, even in complex environments such as the cell membrane.

Our data indicate that mPIEZO1 can occupy a more highly curved state in cell membrane than what has been shown in liposomes and from AF3 predication. Importantly, we revealed, to the best of our knowledge, the first structural insight on blade unbending in the flat state, where they rotate in an anti-clockwise manner to occupy a larger area. We anticipate that the observed unbending might contribute to the gating mechanism, e.g., by stabilizing the channel opening, or acting in extreme stretching condition of the cell membrane. Overall, our data indicate that the blades undergo a spiral rotational motion from a fully extended state in a flat conformation towards a conformation with a small in-plane radius, where the blades highly curve out of the membrane plane. This finding is in line with the predications of AF2 and AF3 and assumption made in an earlier Cryo-EM study^6^ (Fig. 4b).

Further improvements in the labeling strategy, e.g., by simultaneous measurement of the blades with respect to the C-terminal domain of the mPIEZO1 protein will allow one to resolve the curvature of the protein in the membrane. In addition, repeated fast laser-induced vitreous ice melting and reverification can be used to probe conformational changes on the same protein^43^. Furthermore, *in situ* scanning tips can be used to interrogate the response of the protein-membrane system to mechanical stimulation in the cryogenic microscope^44-46^. Findings from spCryo-LM will also be helpful for improving the AF models, which have so far been mainly trained data from isolated proteins. A particularly exciting vision is to realize correlative measurement with Cryo-ET^47,48^, where spCryo-LM provides PIEZO conformational states, while Cryo-ET delivers information on the ultrastructural context of the membrane curvature.

## Acknowledgements

We are grateful to Roderic MacKinnon and Yuxi Zhang for sharing their protocol on cell unroofing on TEM grids. We also warmly thank Ardem Patapoutian and his team for providing the mPIEZO1 plasmid. We would like to thank Franz-Ferdinand Wieser for his crucial contribution to establishing the cryogenic optical microscope and sample transfer pipeline, as well as for his assistance in sketching Figure 1e. We thank Robert Branscheid and Erdmann Spiecker for the loan of the Vitrobot shock-freezing tool in our lab, as well as for assisting in imaging samples using an electron microscope. We thank Simone Ihloff and David Albrecht for help in the cell culture lab, Tobias Utikal and Fabian Greiner for assistance with cryogenics. In addition, we thank Roderic Mackinnon, George Vaisey, Kristian Franze, Franz-Ferdinand Wieser and David Albrecht for valuable comments on our manuscript.

